# Predictive coding and oscillations underlie the optomotor response in distant insect lineages

**DOI:** 10.64898/2026.03.01.708750

**Authors:** Océane Dauzere-Peres, Simon De Wever, Antoine Wystrach

## Abstract

The optomotor response (OMR) refers to animals’ distinctive turning response to rotating visual stimuli. It is assumed to be a predictable Stimulus-Response (S-R) behavior. Here we show in two phylogenetically and ecologically distant insects (ants and earwigs) that it is not the case. Insects’ turning responses were not determined by the detected visual-motion, but by the computation of optic flow prediction errors, which we could infer by manipulating the visual-feedback expected from their own movements. Moreover, both insects displayed regular oscillations of angular velocity, which were modulated by the prediction errors. This produces a closed-loop system yielding complex dynamics and a stochastic moment-to-moment behavior, which can be captured with a simple predictive computational model. We conclude that internal oscillators and predictive coding, as well as their closed-loop interaction, are ancestral features of insect brains conserved across more than 350 million years. The OMR is a by-product of this control system and appears deterministic only when averaging animals’ responses.

## 1 Introduction

Some argue that vertebrate brains are fundamentally predictive coding systems [1–3], but does this apply to insects’ brains too? Insects can generate efference copies – internal neural copies of motor commands – that allow them to predict the sensory consequences of their own actions [4–11]. Notably, work in ants and flies has highlighted the ability of these insects to predict optic flow according to their movements in order to regulate their locomotion [8–11]. How far can this predictive mechanism be generalized during locomotion in insects?

We put this idea to the test using the classical paradigm of the optomotor response (OMR) in two phylogenetically and ecologically distant insect species (earwigs and ants). The OMR is a characteristic orientation response to wide-field visual motion observed in a wide range of animals [12–16] and used to study various aspects of visuo-motor systems [17–24]. Also called ‘optomotor reflex’, the OMR is supposedly based on a highly predictable, simple and dedicated Stimulus-Response (S-R) type of feedforward mechanism [15–17, 25–29], which is not consistent with movement control based on predictive coding.

Our findings show that the OMR is not a deterministic S-R type of behavior. Instead it emerges from the computation of optic flow prediction errors modulating an endogenous oscillator in a closed-loop process. Both ants and earwigs showed the same pattern of results, despite their more than 350 million years of independent evolution and ecology [30]. This indicates that such a predictive-based closed-loop process is a remarkably conserved feature of insects’ brains. A simple neural model of this system captures the results remarkably well.

## 2 Results

### 2.1 Modulation of an oscillator rather than a reflexive response

We placed individual *Cataglyphis velox* ants and *Forficula auricularia* in a virtual reality set-up enabling to decouple the insect’s movements from its visual feedback while precisely recording its locomotion (Fig.1A). When exposed to a rotation of the visual scenery at a constant speed (i.e., a classical OMR paradigm), the turning velocity of the ants and earwigs was, on average, significantly biased in the direction of the external rotation (for ants linear mixed effect model; 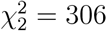, *P* < 0.001, Fig.1B, for earwigs see Fig.S1A,C), showing the expected OMR. Interestingly, for both insects, the angular velocities observed showed a high inter-individual variability, even though they are all a priori in a similar state, and did not match the “correct” speed to compensate for the image rotation, even when averaging at the population level (Fig.1B, Fig.S1A, Fig.S2). This mismatch challenges the idea that the OMR is a stereotyped S-R mechanism dedicated to compensate for external rotations, and prompted us to take a closer look at the underlying dynamics.

**Fig. 1:**
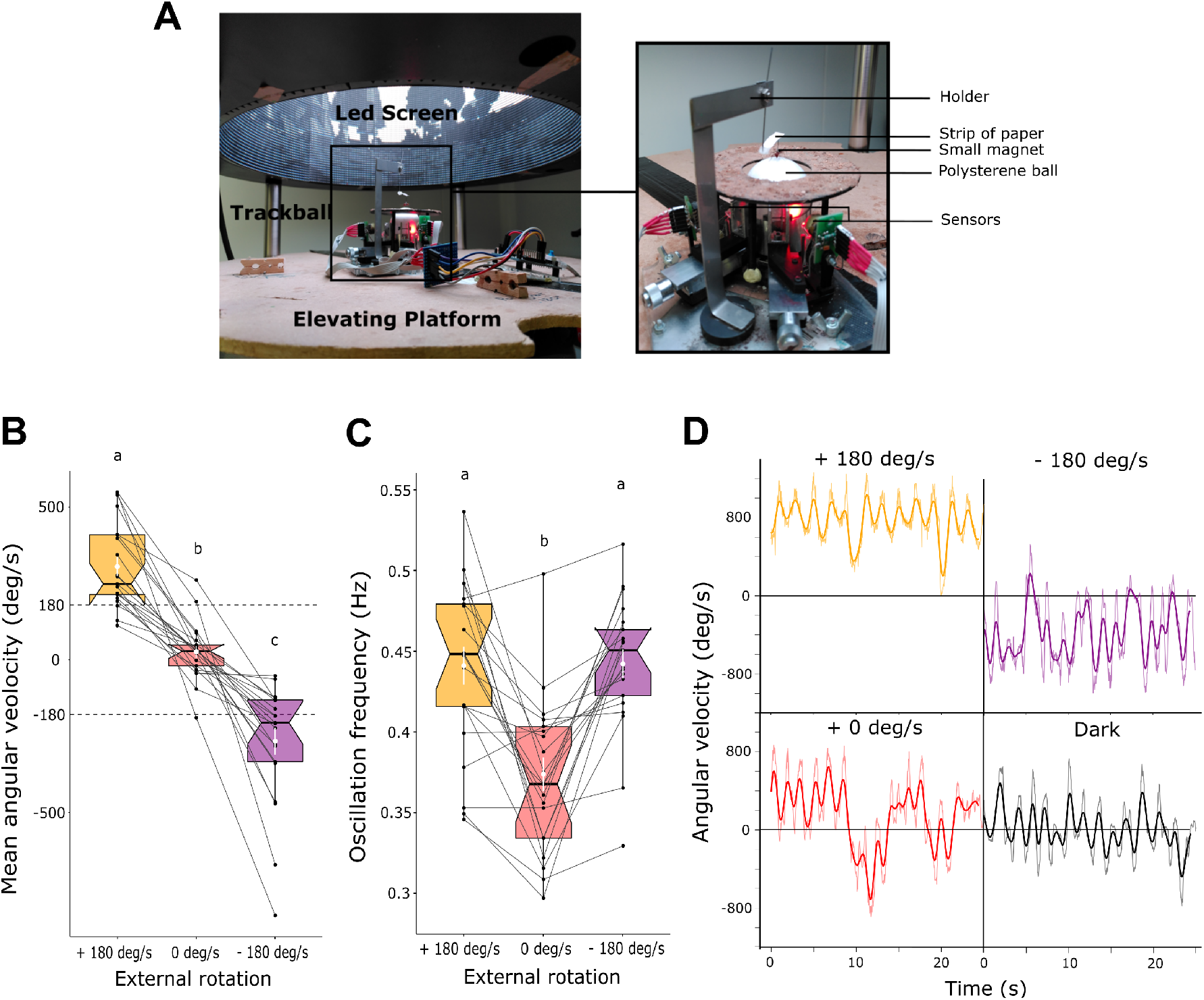
Ants’ optomotor response is a consequence of the modulation of their intrinsic oscillator. **(A)** Virtual reality (VR) experimental set-up. **(B)** Ants’ mean angular velocity (i.e., directional bias) and **(C)** average frequency of ants’ oscillations, depending on the exposure or not to a constant rotating panorama toward the right (−180 deg/s) or the left (+180 deg/s). Data based on 22 ants tested in the VR (n=22 tested in all three conditions). Conditions with different letters show significant differences in post-hoc comparisons; see statistical analysis section. Box plots represent the interquartile range with the median at their center. The white dots and lines represent the means surrounded by their standard errors. Each point corresponds to the response of an individual ant while the lines connect the responses of the same ant across the different conditions. **(D)** Examples of angular velocity signals across time for the different conditions. The dark condition was taken from the periods of 15s between conditions where the LEDs screen was turned off and ants were in the dark. Lighter curves correspond to the raw recordings and the darker ones correspond to the smoothed signals.

During normal locomotion, different insects typically display regular alternations between left and right turns, which results from an internal oscillator located in the insect brain pre-motor area [31–36]. To put in simply, this internal oscillator has a right part and a left part, each making the insect turn in the corresponding direction, constituted of so-called “flip-flop neurons”: the fact that the activity switches from the right to the left neurons generates alternating turning directions. Remarkably, in both species tested here, these oscillations in angular velocity were still conserved during the optomotor conditions, although shifted in the direction of the external rotation, sometimes in the form of regular alternations between turning faster and slower in the same direction (Fig.1D,Fig.S1C). This shows that the insects’ internal oscillator is not silenced during the OMR. The observed shift in oscillations could result from two parallel pathways: one producing the oscillations (the intrinsic oscillator), and one producing a constant turning signal due to the perceived rotation of the scene (i.e., the OMR), summed at the motor output. However, we observed a significant modulation in the frequency of the oscillations in the optomotor conditions compared to when the ants were exposed to a static panorama (linear mixed effect model; 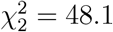, *P* < 0.001, Fig.1C) or when the earwigs were in the dark (Fig.S1B). This would not be expected if the optomotor signal was bypassing the oscillator, shifting the angular velocity signal but leaving its intrinsic rhythm unaltered. Instead, such a frequency modulation is expected if the optomotor signal directly modulates the oscillator by activating one or several of its input regions [36]. We therefore conclude that the OMR results from a dynamic interaction with the activity of the insect’s intrinsic oscillator, rather than from a direct S-R control pathway.

### 2.2 Locomotion under the direct control of prediction errors rather than detected visual motion

We next wondered what was the nature of the signal modulating the oscillator during the OMR. Under the classic S-R perspective, it is generally assumed that the command for turning left or right is based on the *detection* of rotational optic flow (leftwards or right-wards) [14, 16, 17, 25–29]. Alternatively, we reasoned that the OMR could result from a forward model tuned to predict the future sensory outcome of an action, as observed in other contexts and species [1–4, 6, 7, 37, 38]. We know that, at least under certain conditions, insects such as flies [9–11, 27] and ants [8] produce efference copies of their motor commands to cancel the ‘predicted’ part of the optic flow due to their own movements. A difference between the detected and predicted optic flow, so-called prediction error, indicates a displacement of the insect caused by the external environment. Such predictive-coding bears fundamentally different cognitive implications from a S-R mechanism [1–4, 37, 39], although both predict that the animal should, on average, turn in the direction of the external rotation during the OMR.

To disentangle these two hypotheses, we compared the insects’ behavior in two different versions of the optomotor paradigm: open-loop and closed-loop. In open-loop, the animal mounted on the trackball has a fixed orientation and its movements do not modulate the visual stimuli displayed. Conversely, in a closed-loop setting, the animal remains fixed to the trackball but its rotational movements (attempt to turn) are connected to those of the visual display so that the visual motion generated is taken into account, effectively establishing a closed-loop connection with the surrounding scene. Additionally, an external rotation of the scene is consistently applied (Fig.2A). This produces an analogue of the OMR assays where animals are free to physically rotate [26], although fixing the animal as we did here enables a direct comparison with the open-loop condition. The difference between these two versions of the OMR can lead to differences in behavior [40]. Indeed, from the animal’s point of view, the difference is critical.

**Fig. 2:**
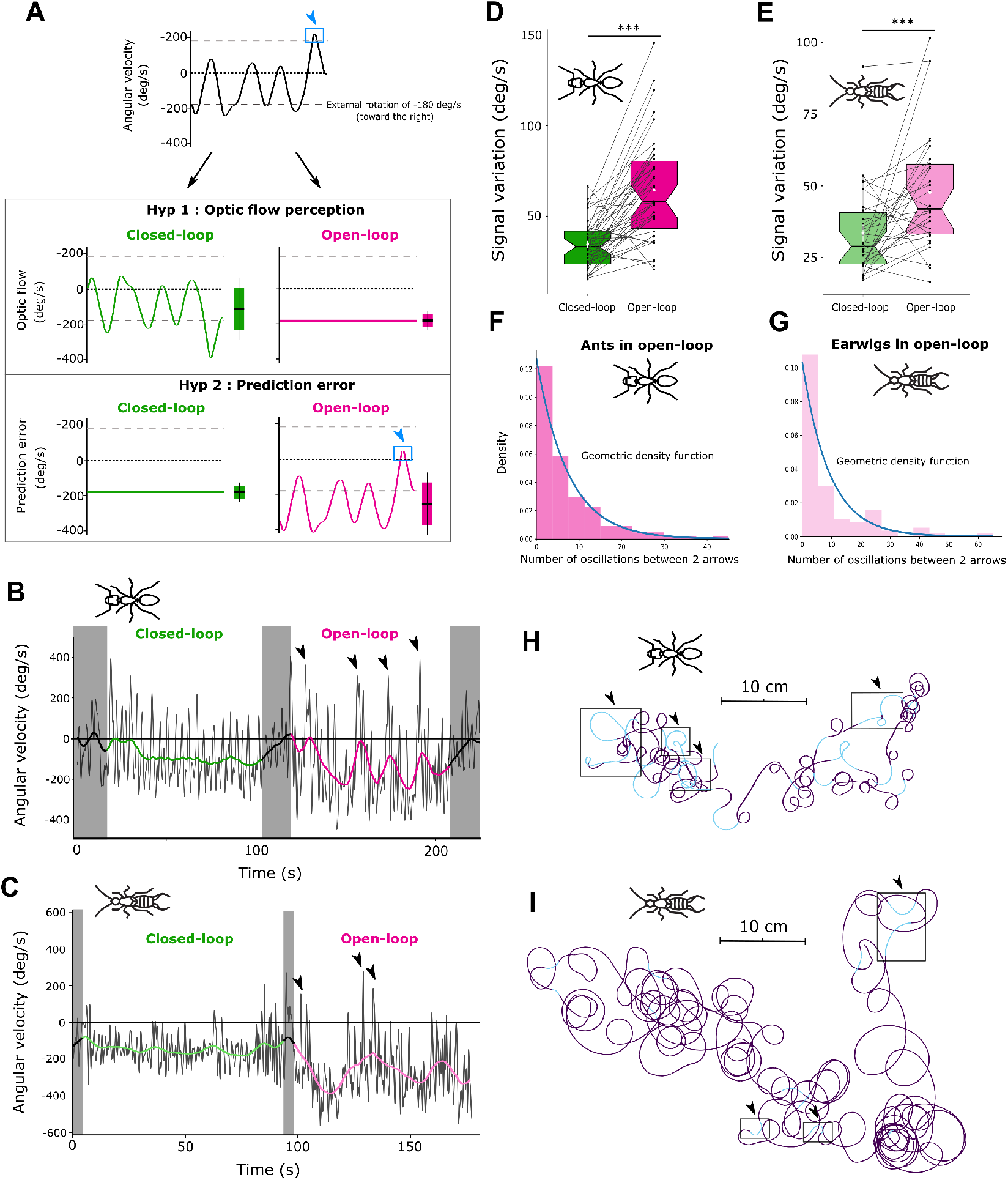
Ants’ optomotor response is driven by optic flow prediction errors. **(A)** Evolution of optic flow perception and prediction error signals given a same theoretical angular velocity signal, either in closed-loop with a constant additional rotation or in open loop with a constant rotation. In both cases the external rotation has the same speed of 180 deg/s (toward the right). The blue arrowheads indicate examples of the ant going against and faster than the rotation, and how it translates into a switch in the sign of the prediction error when in open loop. **(B-C)** Smoothed angular velocity signals of a singular ant (B) or earwig (C) across time (grey curve) and the corresponding signal variation (colored curve) (see method section for more details). Grey boxes and black signal variation curves correspond to the periods between conditions where the LEDs screen was turned off and insects were completely in the dark. In the first condition (green signal variation curve), the animal is in closed-loop with an additional rotation while in the second condition (pink signal variation curve) it is in open-loop exposed to a constant rotation. **(D-E)** Comparison of mean signal variation for ants (D) or for earwigs (E) when the insects performing the optomotor response are in open-loop or in closed-loop with an added rotation. The mean signal variation corresponds to the standard error of the angular velocities in the signal variation curve shown in (B). Data based on 21 ants tested in the VR in both conditions both toward the right and toward the left (n=42 in both conditions) or 30 earwigs tested in the VR in both conditions either toward the right or toward the left (n=30 in both conditions): see Materials and Methods for more details. Box plots represent the interquartile range with the median at their center. The white dots and lines represent the means surrounded by their standard errors. Each point corresponds to the response of an individual insect exposed to an external rotation toward the right or the left. The lines connect the responses of the same insect exposed to a rotation in the same direction. *** = *P* < 0.001; see statistical analysis section. **(F-G)** Density histogram of the number of oscillations between two “arrowhead” events (see method section for how we characterized them), represented in (B) or (C), across all the ants (F) or all the earwigs (G) in open-loop. The blue curve corresponds to the geometric density function fitted with the mean of our data (see method section for more details). **(H-I)** Trajectory corresponding to the angular velocity signal in open-loop shown in (B) or (C). The blue parts correspond to left turns whereas the purple parts correspond to right turns. The arrowheads are matching those in (B).

In the first case (open-loop = external rotation alone), the optic flow on the insect’s retina is caused only by the external rotation of the visual scene and is thus constant. However, the computed prediction error should vary across time with the animal’s movements on the trackball – and can even switch direction (Fig.2A). In the second case (closedloop + external rotation) it is the opposite: the optic flow on the insect’s retina is highly variable – and can switch direction – as it also depends on the insect’s own movements. On the opposite the computed prediction error should remain constant and equal to the added external rotation (Fig.2A). Therefore, if an animal uses the *detected* optic flow to control its behavior (Fig.2A Hyp1), its response should be rather stable in open-loop (i.e., as the *detected* optic flow is constant) but variable in closed-loop condition (i.e., as the *detected* optic flow varies). Conversely, if the animal computes *prediction errors* to control its behavior (Fig.2A Hyp2), its response should be variable when in open-loop (i.e., as the *prediction error* varies), and stable when in the closed-loop condition (as the *prediction error* remains constant).

Our results support the use of prediction errors. The ants’ as well as the earwigs’ oscillations were more stable in the closed-loop condition: the variation of individuals’ angular velocity across time was significantly lower (linear mixed effect model; for ants: 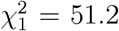, *P* < 0.001, for earwigs: 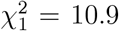, *P* < 0.001, Fig.2B-E) and the variation between individuals’ mean at the population level was also significantly lower (Fig.S3). In contrast, when in open-loop, the insects’ responses varied drastically across time in an apparently chaotic way. Remarkably, in this condition the ants and earwigs even turned frequently at high speeds in the ‘wrong’ direction (arrowheads on Fig.2B-C and Fig.2H-I) even though the optic flow was constant on their retina. These reversal events occurred more frequently in the open loop condition (generalized linear mixed effect model; for ants: 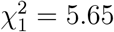, *P* = 0.017, Open-loop: Mean ± SE = 6.71 ± 0.96 %, Closed-loop: Mean ± SE = 3.96 ± 0.79 %, for earwigs: 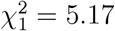, *P* = 0.023, Open-loop: Mean ± SE = 6.44 ± 0.59 %, Closed-loop: Mean ± SE = 4.55 ± 0.52 %) despite that on average both insect species were significantly more biased in the direction of the rotation in this condition (Fig.S3), which is also consistent with the use of prediction errors (Fig.2A). Such anti-optomotor turns, which are not the results of aliasing, have actually been studied and identified in Drosophila [41, 42].

Displaying isolated looping behavior in the ‘wrong’ direction is counter-intuitive. In open loop during which the stimulus is constant, it cannot be explained by a S-R computation, even considering aliasing that occurs at higher external rotation speeds (Fig.S2), and defies the definition of the OMR as a response eliciting a turn in the same direction of a rotating stimulus. However, under a predictive coding perspective, strong reversals in turning direction are expected in open-loop condition. In this condition, the prediction error can reduce and even switch direction as soon as the insect tries to turn faster than the stimuli’s motion (here 180 deg/s or 90 deg/s) in the opposite direction (blue arrows in Fig.2A), thus encouraging the insect to turn further in this direction opposite to the external rotation.

### 2.3 Stochastic rather than predictable

We investigated whether the apparent irregularity of the ants’ and earwigs’ anti-optomotor turns in the open-loop condition followed some predictable pattern across time or was truly stochastic. To do so, we isolated the peaks when ants turned at high speed in the ‘wrong’ direction (opposite to the external visual rotation, see the method section for details) (arrows on Fig.2B-C and Fig.2H-1) – and analyzed the dynamics of their occurrence within each individual. Remarkably, for both insect species the distribution of the number of oscillation cycles separating two consecutive such events fits a geometric distribution, the hallmark of a random process (Fig.2F-G). That is, each event has an equal probability of happening at each time step (each oscillation cycle), regardless of the past of the system, and thus shows the stochastic nature of the behavior emerging from this system.

### 2.4 Continuous closed-loop control

Considering prediction errors together with stochasticity and intrinsic oscillators working as a closed-loop process (Fig.3B) provides a straightforward explanation for why ants (Fig.1D, +0 deg/s) mounted in open-loop in front of a static panorama (i.e., that is, without external rotation or reafference) also display highly irregular, shifted oscillations [8]. In this condition, looping behaviors identical to the one observed in the OMR – but emerging here from a still visual input – are expected due to the mismatch between the predicted presence and detected absence of optic flow, which generates strong prediction error signals. Random changes in the direction of these looping behaviors toward the left or the right further highlight the stochastic process acting with the internal oscillator. Such a ‘self-induced’ looping behavior shows that the OMR does not result from a mechanism dedicated to respond to external movements of the scenery (Fig.3A), but emerges from closed-loop mechanisms operating continuously to control the insect’s movements.

**Fig. 3:**
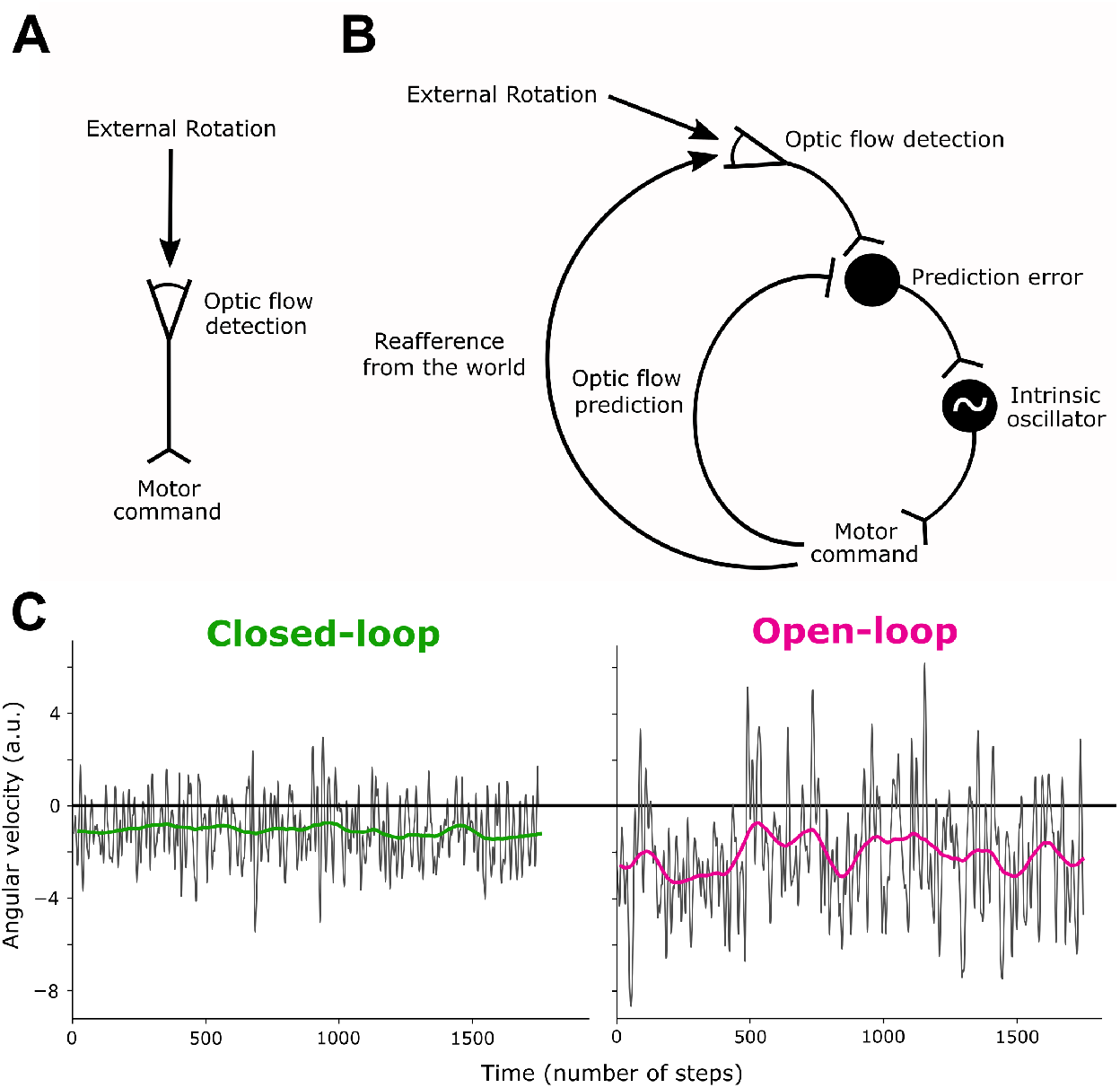
Conceptual models underlying the optomotor response. **(A)** The optomotor response explained as a dedicated stimulus-response mechanism. This model produces a response only in the presence of optic flow. The direction of the optic flow dictates the direction of the motor response. **(B)** Model of ants’ visuo-motor control operating as a continuous, closed-loop process, involving an endogenous oscillator and the production of prediction errors. This model generates the optomotor response as a by-product. The direction of the motor response results from non-linear interactions and varies through time, with or without optic flow. **(C)** Examples of simulations of the model (B) in closed-loop and open-loop (by removing the ‘reafference’ feedback) with an external rotation set to −2 (arbitrary angular velocity scale). Smoothed angular velocity signals across time (thin curve) and the corresponding signal variation (thick curve) are reported in arbitrary units (a.u.). See Materials and Methods, supplementary text and Table.S1-S2 for more details on the algorithm used.

As a proof of concept, we implemented a simple neural model of the suggested closedloop control system. We used optic flow prediction based on motor information to compute prediction errors, with the latter modulating an endogenous neural oscillator (Fig.3B). Given the addition of noise to the activity of the intrinsic oscillator to account for neural firing stochasticity and unknown input, the model reproduces the complex dynamics found in open-loop and closed-loop in both insects impressively well (Fig.3C), and offers a plausible neural implementation of this control system. The fact that monocular ants (with one eye covered by paint) displayed an optomotor response approximately half as strong (Fig.S4), supports the idea that each eye computes their own prediction errors, which have an additive effect on the oscillator [8].

## 3 Discussion

Our results invite us to reconsider the nature of the mechanism underlying the OMR: it is not a stereotyped, predictable behavior, nor a dedicated S-R or reflex mechanism, as suggested in drosophila for which the anti-optomotor turns also seem inconsistent with the idea [41]. Instead, the OMR is a by-product of interacting mechanisms – involving an intrinsic oscillator [8, 31], inherent stochasticity, and crucially, the computation of prediction errors. This system operates as a closed-loop process to continuously control the movements produced by the insect itself, with or without external rotation of the scenery (Fig.3). Remarkably, this visuo-motor control system is very similar in ants and earwigs, despite their remote ancestry and different ecologies, indicating that it may have been conserved for at least 350 million years [30]. More generally, it revealed that predictive coding was likely already present in ancient insect brains or, in the case of convergence, that it is important enough to appear separately in two very distant species.

### 3.1 Predictive coding as a framework for understanding behaviors?

Predictive coding, notably through efference copies generating prediction errors, is increasingly viewed as a critical concept to understand vertebrates’ brains [1–3], notably regarding the optokinetic system [43–45]. In insects, efference copies of motor commands have been shown in several contexts [5–7, 38], including the inhibition of optic flow responsive neurons in visual areas [9–11, 27]. Efference copies in insects might target only specific sensory neurons [11], operate only in some contexts [27], and may not necessarily mediate complex predictive 3D representations such as the forward models involved to control body movements in humans [4, 37, 46]. Nonetheless, the fact that a spontaneous and ancestral visuo-motor behavior such as the OMR is based on prediction errors as shown here, together with the recent realization that there is a massive amount of neural feedbacks from motor areas in insect brains [47–50], suggests that closed-loop control and predictive coding should not be treated as exceptional, but rather as a general framework to understand the insects’ brains too. Consequently, neural recordings of fixed animals, even in sensory areas, should not automatically be interpreted as *detection* signals since they may well be *prediction error* signals, thus sensitive not only to the stimuli but also to the motor behavior, which is often ignored and would be needed for correct interpretation of the signals.

### 3.2 Stochasticity and oscillations to explore various regimes of activity?

Inherent stochasticity in brains arises from the noise underlying the probabilistic spiking time of neurons [51]. Studies in humans and primates show that this noise might actually be advantageous for decision-making, notably by preventing deadlocks and exploring other regimes in the system [39]. The stochastic behavior observed here in ants and earwigs embodies this idea remarkably well, but in the context of a basic locomotor behavior rather than a cognitive function. In open-loop, and despite the strong sensory evidence that the world is spinning in one direction, noise – on top of the continuous production of intrinsic oscillations – enables the insects (Fig.2), and our model (Fig.3C) to escape the strong attractor of turning in this direction and explore other regimes, such as occasionally turning in the anti-optomotor direction, given a random probability (Fig.2D-E). Intrinsic neural oscillators are ancestral and widespread in invertebrates and vertebrates, and the ‘blind’ exploration of the world they embody is useful in the context of navigation [31, 33, 52]. Perhaps more generally, the universal presence of oscillations in neural activity [53, 54] together with noise, is key for escaping and exploring various regimes of brain activity.

### 3.3 Conclusion

Some have argued that neuroscience suffers from a detrimental, reductionist bias resulting partly from methodological constraints, notably the need to fix the animal to record its neural activity [40, 55], and the lack of consideration for behavior itself [56]. In the case of the OMR, it seems indeed that the need to simplify the complexity of natural behaviors has yielded a parallel and tacit simplification of our theoretical understanding of the underlying mechanisms. Simplification through the use of open-loop design to record neural activity [28, 57–59], or averaging the animal’s response across time and/or individuals [18, 19, 21, 23], buys predictability but hides the fundamentally complex dynamics at play [60].

## 4 Materials and Methods

### 4.1 Studied species and housing conditions

Colonies of *Cataglyphis velox* were collected near Seville (Spain) in 2020 and transferred to the CRCA in Toulouse (France). Colonies consisted of one or more queens, workers (measuring up to 12 mm [61]), and brood. They were housed in vertical nests made of aerated concrete with manually dug galleries and chambers. These nests were kept in darkness in a climate-controlled room (24–30 °C, 15–40% RH). Each nest was connected via a transparent tube to a 40 × 30 cm sand-covered foraging arena. The arena was exposed to a 12:12 h light/dark cycle under a heating lamp. Ants had ad libitum access to water and a 40% sucrose solution, and were provided mealworm pieces three times per week.

European earwigs (*Forficula auricularia*), are subsocial dermapterans measuring 13–14 mm excluding forceps [62]. They were collected in June 2024 from a peach orchard in Saint-Marcel-lès-Valence (France). Individuals were maintained at the IRBI in Tours (France) in plastic containers containing moist sand as substrate and an egg carton as nests. They were supplied with standard laboratory food ad libitum [63]. Containers were kept under a 12:12 h light/dark cycle at 18–20 °C. In July 2024, the individuals were transferred to the CRCA in Toulouse, for experimental studies.

### 4.2 Experimental set-up: Virtual world and data acquisition

The virtual reality (VR) set-up consisted of a trackball system surrounded by a 360° cylindrical LED display (Fig. 1B). Insects were tethered to walk on a polystyrene ball floating on an air cushion, compensating their movements [64]. The ants were marked with a small dot of magnetic paint on their thorax and mounted in a fixed orientation using a holder with a micro-magnet. The earwigs were mounted in the same way, with the paint applied to the elytra. The trackball was placed on a lifting platform, enabling to raise the mounted insects within the VR screen. The cylindrical LED display (50 cm diameter, 76 cm height) contained 73,728 RGB LEDs, yielding a resolution of 0.94°/pixel, which is higher than the resolution of *Cataglyphis* ants’ eyes (> 2 deg/pixel [65]). The scenery projected on the screen was controlled online by a computer using custom codes implemented in a Unity® 2020.1 environment, and updated online according to the insect’s movements, which were recorded by two sensors tracking the ball. A camera placed on the top of the VR set-up allows us to monitor that the insect is attached properly during experiments.

In all experiments, the display consisted of a black-and-white panoramic scene mimicking the natural habitat of *C. velox*, thus providing realistic rotational optic flow. Translational optic flow was not simulated. The VR enabled us to manipulate the relationship between the rotational movements performed by the insects and the visual rotation of the scenery. The VR system was run in two modes: (1) open-loop, in which insects’ movements had no effect on the panorama (which was either static or rotating at a constant speed toward the left or the right), and (2) closed-loop, in which the insect’s movements controlled the rotation of the panorama (gain of 2 for ants, 1 for earwigs) while an additional constant external rotation was applied independently of the insect’s behavior.

### 4.3 Experimental protocol

Tested ants were randomly picked from the foraging area and marked individually using a color code composed of a dot of paint on their thorax and two dots on their abdomen. Magnetic paint was also applied to the middle of their thorax to enable them to mount the trackball. The painted ants were then placed for a few minutes in a small pot to let the paint dry, and were mounted on the trackball with the platform in the lowest position. The screen of the VR was first turned off while the platform was raised to immerse the ant in the VR set-up; the screen was then switched on and the trackball movements were recorded. Once the trial was over, the ant was placed back in its nest.

Tested earwigs were randomly picked from a terrarium in Tours and brought to the laboratory in Toulouse in petri dishes the day before the experiment. Magnetic paint was applied to the elytra of the chosen individuals. The painted earwigs were then also placed for a few minutes in a small pot to let the paint dry before being mounted on the trackball with the platform in the lowest position. Once the trial was over the earwigs were placed back in separate petri dishes.

For the first series of experiments with ants, the data presented above correspond only to a portion of the data collected. For the purpose of this study, we only analyzed ants that had both their eyes uncovered in the main text; however, in the entire experimental protocol, an additional dot of paint was also either applied on their left eye (LE), right eye (RE), or for the group we used here, above their left eye (2E). The data on monocular ants were analyzed in Fig.S4. At the end of the trial, the paint on their head was removed before releasing the ants back in their foraging arena. The individually marked ants could be picked up from the nest multiple times to be tested in the different conditions (LE, RE, and 2E) in a randomized order. In total, 22 different ants with both their eyes uncovered were tested in open-loop. The ants were tested successively without leaving the VR under three conditions: with a constant external rotation of 180 deg/s toward the left, 180 deg/s toward the right, and 0 deg/s (static panorama). The order of the two optomotor conditions was randomized, but the static condition was always tested last. Each condition lasted for 180 seconds and the screen was turned off for 15 seconds between two conditions (reproducing a dark condition).

For the second series of experiments with ants, we tested 21 ants in optomotor condition both in open-loop and closed-loop. The ants were tested successively without leaving the VR under 4 different conditions in a pseudo-randomized order: with a constant external rotation of 180 deg/s toward the right or toward the left (open-loop), and in closed-loop with an additional constant external rotation of 180 deg/s toward the right or the left. Each condition lasted for 90 seconds and the screen was turned off for 15 seconds between two conditions. The speed of the external rotation of 180 deg/s for both experiments was chosen according to preliminary experiments to maximize ants’ response (see Fig.S2).

For the experiment with earwigs, we tested 15 males and 15 females in the dark and in optomotor conditions in closed-loop as well as open-loop. The earwigs were tested successively without leaving the VR under 3 different conditions: the first condition was always in the dark (LED screen turned off), then, they were exposed in a random order to a constant external rotation of 90 deg/s (open-loop) and in closed-loop with an additional constant external rotation of 90 deg/s. Half the males and half the females were tested with the rotation going toward the right, and the other halves with the rotation toward the left. Each condition lasted 90 seconds and the screen was turned off for a few seconds between the two optomotor conditions.

In closed-loop the gain of the ant controlling the rotation of the panorama was set to 2 (instead of 1) according to previous experiments showing that this generated optic flow closer to their expectation [8]. In the absence of such information on earwigs, and because they are heavier and stronger than ants, this gain was left at 1. If the gain does not correspond exactly to ants’ expectation, it could leave residual unpredicted optic flow, however, if present, these prediction errors should be much smaller (and thus way less impactful) than when ants are in open-loop.

### 4.4 Data transformation

All statistical analyses were done using R v.4.0.3 [66], whereas the transformation of the raw data acquired by the system into exploitable variables and their visualization were done with Python v.3.9.7.

For all the trials of each insect, the raw movement detection data collected was split between the different tested conditions and transformed into meaningful variables on ants’ locomotion using a Python code. That way, the virtual trajectory as well as the angular velocity over time (negative when going right and positive when going left) could be extracted. To decrease the natural noise of the recording system (recording at Mean ± SD = 45.3 ± 1.9 fps), the raw angular velocities were resampled to have a data point every 33 ms (30 fps) and smoothed by running twice an average filter with a sliding window of 20 frames long for ants or 15 frames long for earwigs. The length of the window was chosen so that it was shorter than the length of an oscillation and produced clean signals, effectively removing the noise of the recording system. The length of the window was shorter for earwigs as they produced oscillations with a higher frequency. This smoothed signal was then filtered to delete periods during which the insects stopped walking by removing data points between 3 and −3 deg/s for more than 20 consecutive frames. On this smoothed and filtered signal, we detected oscillations by extracting local extremums such as one oscillation corresponded to the succession of three extremums, or in other words, the alternation between an increase phase and a decrease phase (no matter the order), each with a minimum amplitude of 5 deg/s. We chose this method rather than a Fourier analysis because this enables to average the length of the oscillations in different regimes, and not select the one most present.

For the second series of experiments with ants and for the earwigs experiment, we also produced another highly smoothed curve corresponding to the average variation of the signal by running twice an average filter with a sliding window of 200 frames long on the filtered signal. To quantify the mean signal variation we took the standard deviation of the angular velocities in this very smoothed curve. In addition, the extremums going against the external rotation (minimums with a left external rotation and maximums with a right external rotation) were also extracted, for all the oscillations detected in the angular velocity signals. We could then quantify for each insect the proportion of those extremums either in open-loop or closed-loop conditions that were in the highest 5% overall. In addition, when insects were in open-loop, we calculated the intervals by counting the number of oscillations between two of those extremums in the top 5% of the optomotor conditions. We wanted to see if those high extremums happened truly randomly, and to do so plotted a density histogram of those intervals for both earwigs and ants. In a geometric law, a success has an equal probability (p) of happening at each time step (here each oscillation). The geometric distribution then gives the probability that there are X failures (oscillations with lower extremums) before the first success (oscillation with a higher extremum). To see if high extremums happened randomly with the same probability p at each time step, we thus plotted the probability mass function of the geometric law with the parameter p fitted according to the mean of our intervals (p=1/(mean+1)). We were then able to compare our data distributions and the matching geometric distributions.

### 4.5 Statistical analysis

Two continuous response variables were computed for each condition and in each trial: the average angular velocity, corresponding to the overall directional bias in the trajectory of the insect, and the frequency of insects’ oscillations, calculated as the number of oscillations divided by trial duration (excluding inactive periods). For the second series of experiments with ants and for the earwig experiments, we also quantified signal variation as the standard deviation of the very smoothed curve.

All analyses were conducted in R using linear mixed-effects models (LMMs) or generalized linear mixed-effects models (GLMMs) with the package *lme4* [67]. In all models, insect identity was included as a random factor to account for repeated measurements. Model assumptions were checked visually by inspecting QQ plots (normality of residuals) and residuals versus fitted values (homoscedasticity). P-values were obtained with type II Wald chi-squared tests and compared to a significance level of 0.05. When effects involving more than 2 groups were significant, post-hoc pairwise comparisons were performed using the same model on relevant data subsets, and P-values were adjusted with Holm–Bonferroni correction [68].

For the first series of experiments with ants, we used LMMs to test for the effect of the direction of the external rotation (factor with 3 levels: right, left and none (static panorama)) on the average angular velocity of the ants as well as the frequency of their oscillations. For the average angular velocities, we used the signed square-root transformation (square-root transformation on the absolute value multiplied by the sign of the original value) since the values could be negative (when biased toward the right). To check that the effect we found on the frequency of the oscillations was truly caused by the visual condition and not the sequence of the conditions (since the static panorama was always presented last) we verified that there was no effect of time by looking within the conditions. To do so we divided each condition into periods of 15 seconds and used the same LMM with the order of the periods within each condition (discrete variable) as an additional variable. Indeed, time had no significant effect on the frequency of the oscillations (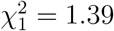, *P* = 0.239), while the external rotation was still significant (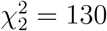, *P* < 0.001).

For the second series of experiments with ants, we used a LMM to test for the effect of the set-up condition (factor with 2 levels: open-loop (gain 0) and closed-loop (gain 2)) on the signal variation. In addition, we used a GLMM for proportional data with the binomial distribution and the link function logit to compare the proportions of the top 5% highest anti-optomotor extremums performed by each ant for each optomotor direction happening when they were in open-loop versus closed-loop (factor with 2 levels). We checked for overdispersion using the function check overdispersion of the R package performance [69], and since overdispersion was detected, we added observation-level random effects to deal with it [70]. In both models, the direction of the rotation (factor with two levels: left or right) was initially put into the model to check that there was not an unexpected asymmetry, but it was dropped after it was not found to be significant.

For the experiment with earwigs, we carried out the same analysis than with the second series of experiments with ants: we also used a LMM to test for the effect of the set-up condition (factor with 2 levels: open-loop (gain 0) and closed-loop (gain 1)) on the signal variation, and a GLMM for proportional data with the binomial distribution and the link function logit to compare the proportions of the top 5% highest anti-optomotor extremums of each earwig happening in open-loop versus closed-loop (factor with 2 levels). We checked for overdispersion using the function check overdispersion of the R package performance [69], and no overdispersion was detected. In both models, the direction of the rotation (factor with two levels: left or right) and the sex of the individual (factor with two levels: male or female) were initially put into the model but they were dropped after they were not found to be significant.

### 4.6 Neural model

We implemented the simple toy model presented in Fig.3B using Python to compare the predicted dynamics with those observed in insects in open-loop or closed-loop conditions. Neurons were modeled as simple linear integrators, that is, performing a weighted sum of their input signals, the weights indicating the strength of the input connections. The sensory neuron takes the value of the optic flow detected (see below). The prediction error neuron activity corresponds to the difference between the sensory neuron, by which it is excited, and the prediction (see below) by which it is inhibited. The prediction error neuron projects to two mutually inhibiting motor neurons (left and right), forming an intrinsic oscillator. The angular velocity is updated using the difference between the activity of the left and the right motor neurons of the oscillator. The optic flow detected by the sensory neuron corresponds to the angular velocity (reafference) plus the constant value of the external rotation (exafference). Note that in open-loop condition (i.e., in the absence of reafference), the optic flow detected corresponds only to the external rotation, and is thus constant. See supplementary text for the detailed algorithm and Table S1-S2 for the parametrization used. Note that the values of the parameters used have little importance for the use we have of the model, we are looking at qualitative differences according to the optomotor assay (open or closed-loop), whereas the other parameters influence the output quantitatively identically in both cases.

In Fig.3C, the angular velocity signals obtained with this model were smoothed by running twice an average filter with a sliding window of 5 frames long, and the signal variation (thick line) was obtained by running twice an average filter with a sliding window of 150 frames long.

## Acknowledgments

We thank Paul Graham and Filipe Pinto-Teixeira for their helpful comments on the manuscript, as well as Jeanne Legendre who participated in the data collection.

## Funding

Funder: European Research Council

Grant reference number: EMERG-ANT 759817

Author: Antoine Wystrach

## Authors’ contribution

O.D.P. Conception and Design of experiment, Data collection, Analysis and interpretation of data, Drafting and revising the article.

A.W. Conception and Design of experiment, Analysis and interpretation of data, Drafting and revising the article, Supervision of project.

S.D.W. Data collection, Revising the article.

## Data and code availability

The processed and raw data supporting the figures as well as the codes used to process the raw data will be made available in a GitHub repository (https://github.com/antnavteam) upon peer-reviewed publication.

## Supplementary information

### Supporting Information Text

Algorithm used to model insects’ visuo-motor system (Fig.3B)

Initialization:

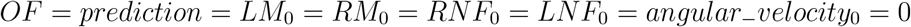

For each time step:

1. Computation of the optic flow, the predicted optic flow and the resulting prediction error

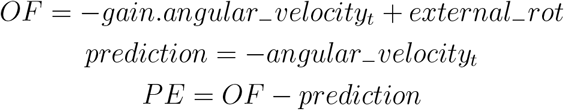
2. Impact of the prediction error on the oscillator

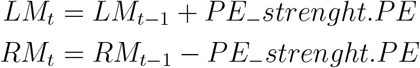
3. Intrinsic oscillator dynamic

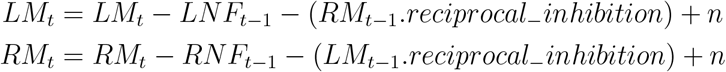

with *n* ↪ *N*(0, *noise*)

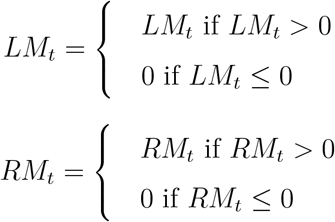

Update the internal negative feedback for the left and right motor neuron:

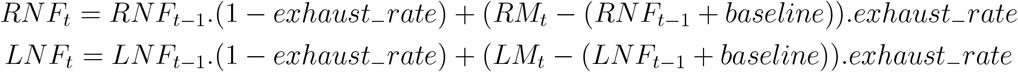
4. Computation of the angular velocity

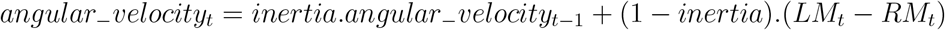

**Table. S1.** Variables used in the algorithm to model insects’ visuo-motor system (Supporting information text)

| Variable | Description |
| --- | --- |
| **Neurons** |  |
| OF | Optic flow detector neuron |
| PE | Prediction error neuron |
| LM | Left motor neuron of the oscillator |
| RM | Right motor neuron of the oscillator |
| prediction | Prediction neuron (predicted optic flow in arbitrary units) |
| **Other variables** |  |
| angular velocity | Angular velocity of the agent (arbitrary units) |
| RNF | Right motor neuron negative internal feedback |
| LNF | Left motor neuron negative internal feedback |

**Table. S2.** Parameters used in the algorithm to model insects’ visuo-motor system (Supporting information text)

| Parameter | Description | Value |
| --- | --- | --- |
| **Fixed parameters** |  |  |
| inertia | Conservation of the current angular velocity between each time step | 0.7 |
| baseline | Firing rate of the oscillator motor neurons without any external modulation | 0.5 |
| reciprocal inhibition | Synaptic weight of the reciprocal inhibition between the two neurons of the oscillator | 0.9 |
| exhaust rate | Rate at which the negative internal feedback is modulated between each time step | 0.15 |
| noise | Variance of the normal distribution used to sample noise values added to the activity of the oscillator's motor neurons | 1.5 |
| PE strength | Weight of the prediction errors on the activity of the oscillator neurons | 0.25 |
| **Free parameters** |  |  |
| gain | Relation between the angular velocity and the optic flow detected | 0 (open-loop)<br>1 (closed-loop) |
| external rot | Additional optic flow which is not generated by the angular velocity (arbitrary units) | - |

**Fig. S1:**
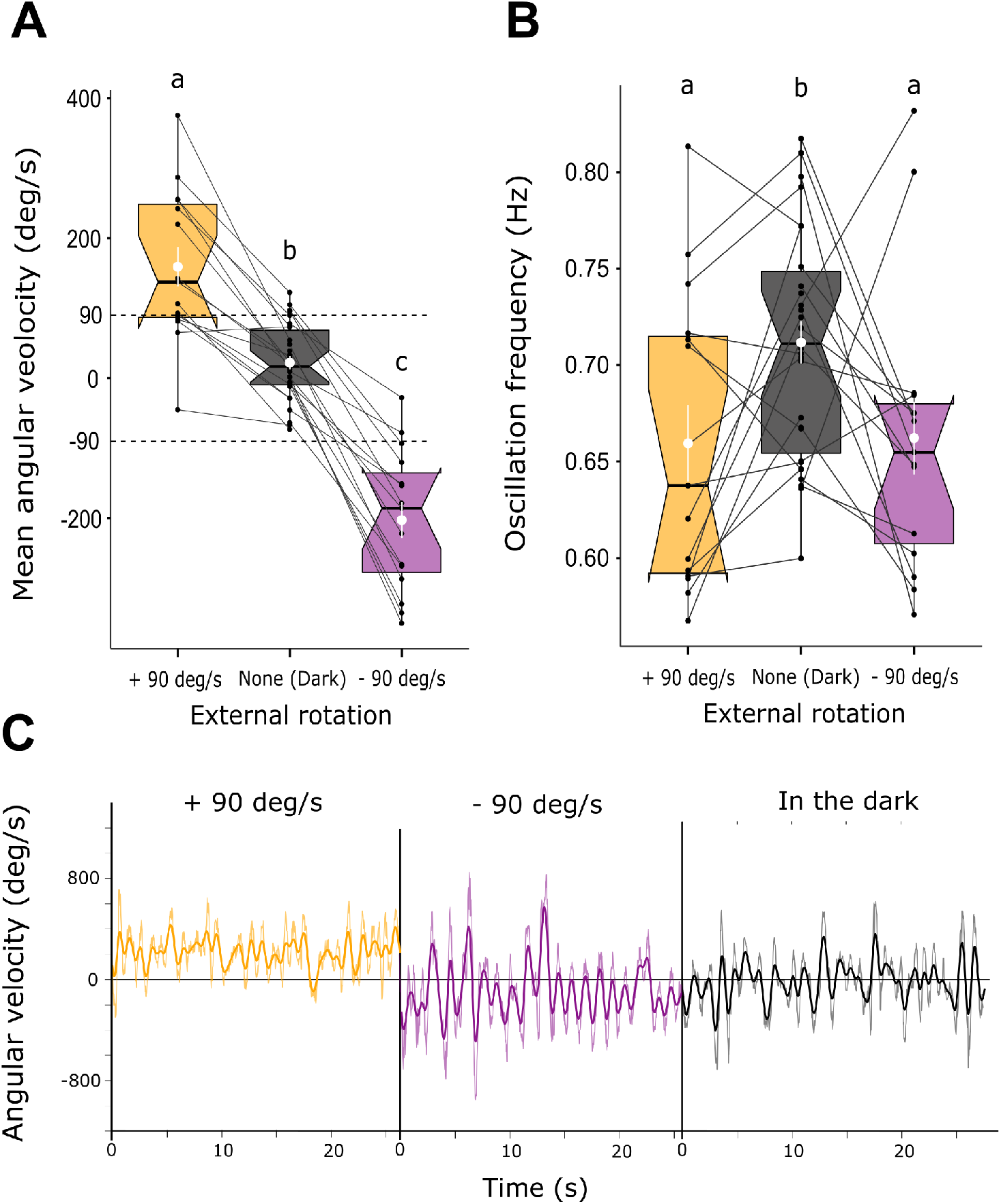
Earwigs’ optomotor response is also a consequence of the modulation of their intrinsic oscillator. **(A)** The earwigs’ mean angular velocity (i.e., directional bias) and **(B)** average frequency of their oscillations, depending on the exposure or not to a constant rotating panorama toward the right (−90 deg/s) or the left (+90 deg/s), see Materials and Methods for more details on the experiment. We used LMMs to test for the effect of the external rotation in open-loop (factor with 3 levels: external rotation toward the right, toward the left or no external rotation (in the dark)) on both behavioral parameters. The identity of the individual was included as a random factor to account for repeated measurements. The sex of the individual (factor with two levels: male or female) and its interaction with the exposure of the rotation were initially included in the models but were dropped after their effects were not found to be significant. There was a significant effect of the external rotation on the earwigs’ mean angular velocity (linear mixed effect model; 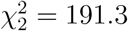, *P* < 0.001) and on the frequency of their oscillations (linear mixed effect model; 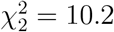, *P* = 0.006). To check that the effect we found on the frequency of the oscillations was truly caused by the visual condition and not the sequence of the conditions (since the dark was always the first condition) we verified that there was no effect of time by looking within the conditions. To do so we divided each condition in periods of 15 seconds and used the same LMM with the order of the periods within each condition (discrete variable) as an additional variable. Time had no significant effect on the frequency of the oscillations (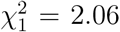, *P* = 0.151), while the significance of the external rotation remained ( 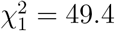 = 49.4, *P* < 0.001). Box plots represent the interquartile range with the median at their center. The white dots and lines represent the means surrounded by their standard errors. Each point corresponds to the response of an individual earwig while the lines connect the responses of the same earwig across the different conditions. **(C)** Examples of angular velocity signals across time for the different conditions

**Fig. S2:**
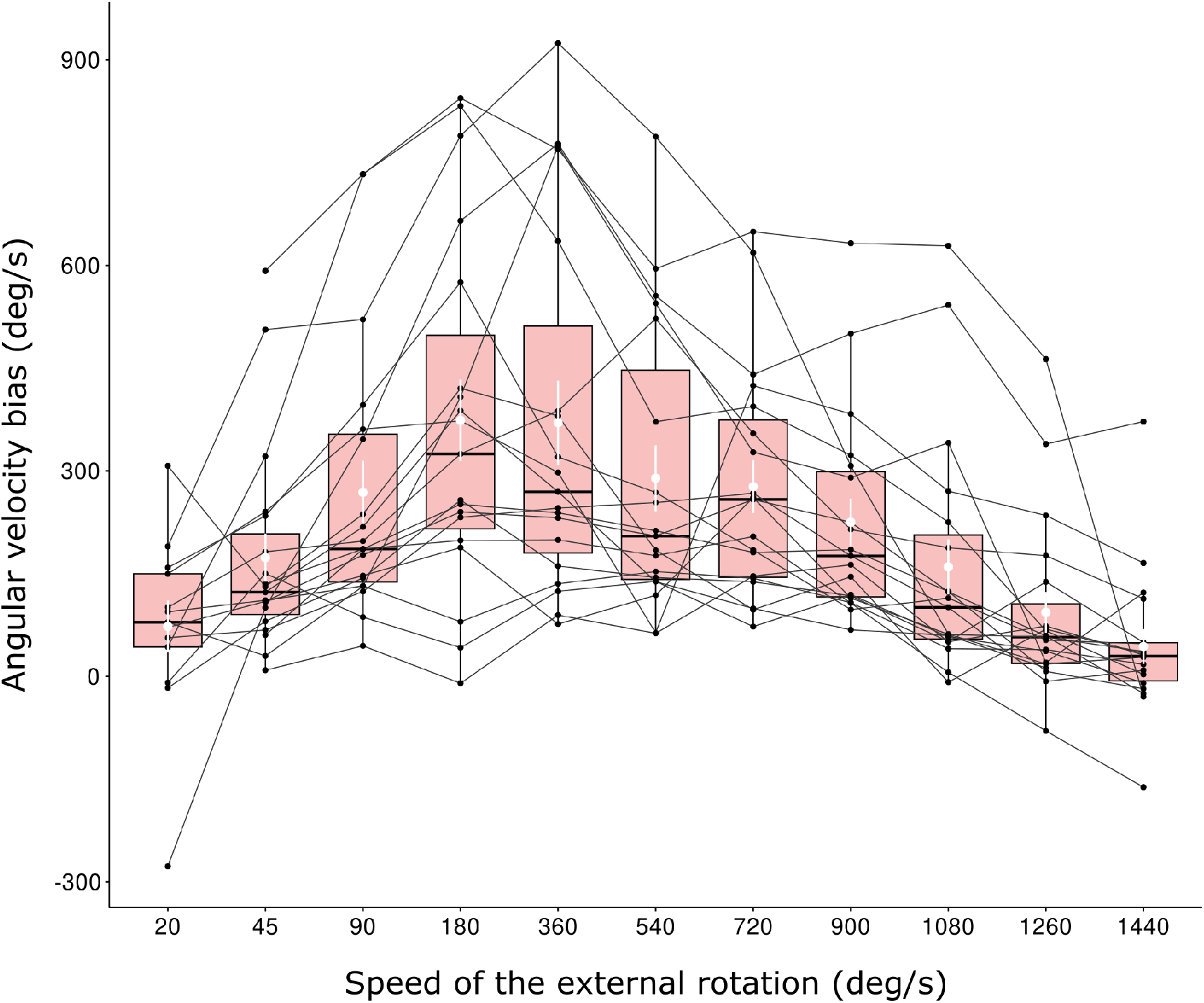
Sensitivity of ants’ optomotor response to different rotational speeds in open-loop. Evolution of the ants’ average angular velocity bias in the direction of the external rotation (i.e., rotation bias) for different rotational speeds in open-loop. These data were collected by placing 19 ants in the VR and exposing them successively to 11 different rotational speeds of the panorama for 60 seconds, going from 20 deg/s to 1440 deg/s (except for 6 ants that were not tested with a speed of 20 deg/s (n=13) and 1 ant which was not tested with a speed of 1440 deg/s (n=18)). Half of the ants were tested with a rotation toward the right (n=9) and the other half with a rotation toward the left (n=10). In both groups, half of the ants were tested with increasing speeds (from 20 to 1440 deg/s, n=4 in the rightward rotation group and n=5 in the leftward rotation group) and the other half with decreasing speeds (from 1440 to 20 deg/s, n=5 in the rightward rotation group and n=5 in the leftward rotation group). Box plots represent the interquartile range with the median at their center. The white dots and lines represent the means surrounded by their standard errors. Each point corresponds to the bias of an individual ant while the lines connect the responses of the same ant across the different speeds.

**Fig. S3:**
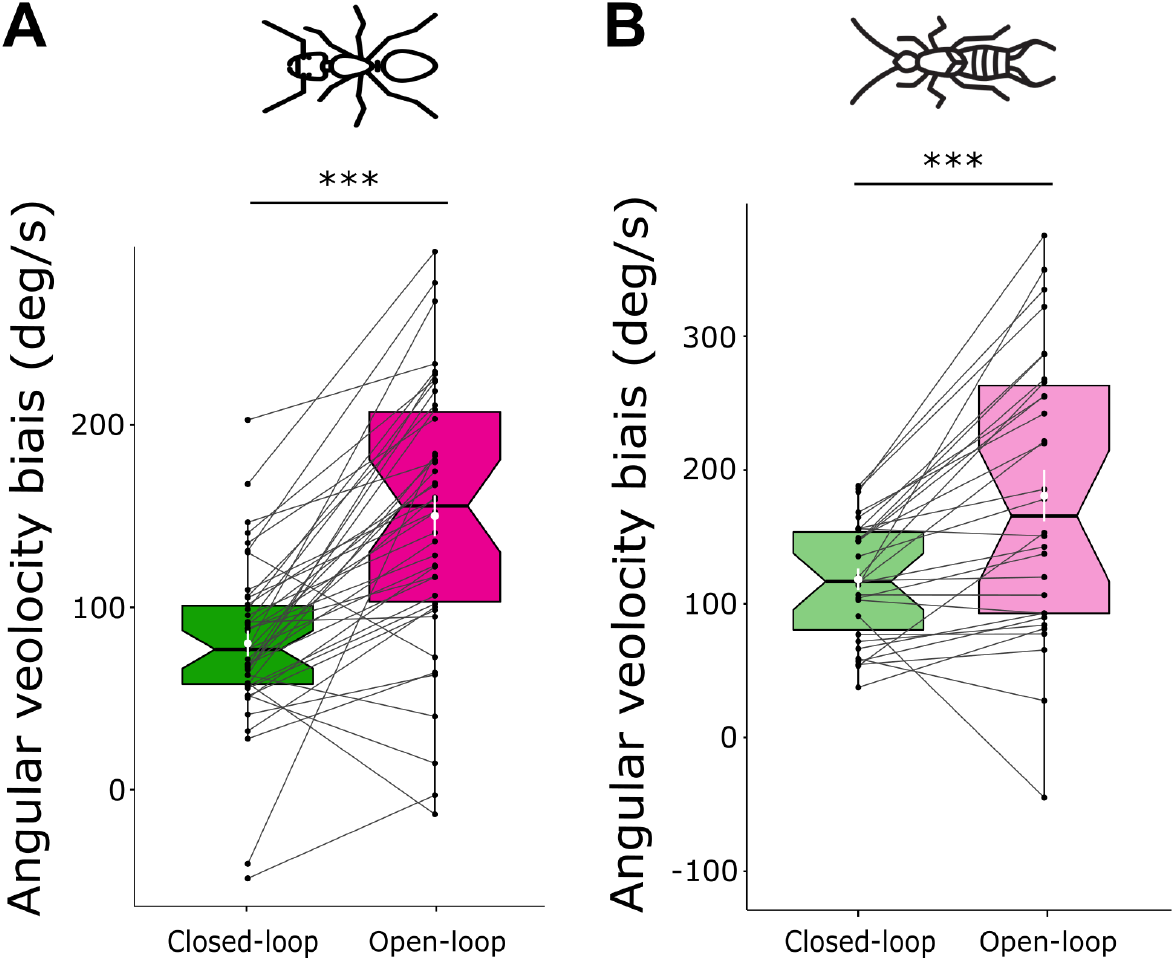
Ants and earwigs directional bias in optomotor condition in closed-loop versus open-loop. **(A)** Comparison of the average angular velocity bias in the direction of the external rotation (i.e., optomotor bias) between the ants performing the optomotor response in open-loop versus in closed-loop with an added rotation. Data based on 21 ants tested in the VR in the two conditions both toward the right and toward the left (n=42 in each conditions), see Materials and Methods for more details on the experiment. We used a LMM to test for the effect of the set-up condition (factor with 2 levels: open-loop and closed-loop) on the angular velocity bias. The identity of the individual was included as a random factor to account for repeated measurements and the direction of the rotation (factor with two levels: left or right) was initially put into the model as a fixed factor but it was dropped after its effect was not found significant. In open-loop the ants were significantly more biased in the optomotor direction than in closed-loop (linear mixed effect model; 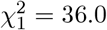, *P* < 0.001) and the variance of their responses was also significantly more important in open-loop than in closed-loop (Levene test on the variance: *F* = 9.58, *P* = 0.003). **(B)** Comparison of the average angular velocity bias in the direction of the external rotation (i.e., optomotor bias) between the earwigs performing the optomotor response in open-loop versus in closed-loop with an added rotation. Data based on 30 earwigs tested in the VR in both optomotor conditions (open-loop and closed-loop). The earwigs were either tested with an external rotation toward the right or toward the left: see Materials and Methods for more details on the experiment. We used a LMM to test for the effect of the set-up condition (factor with 2 levels: open-loop or closed-loop) on the angular velocity bias. The identity of the individual was included as a random factor to account for repeated measurements. The sex of the individual (factor with two levels: male or female), its interaction with the set-up condition, and the direction of the rotation (factor with two levels: left or right) were initially included in the model but they were dropped after their effects were not found significant. In open-loop the earwigs were significantly more biased in the optomotor direction than in closed-loop (linear mixed effect model; 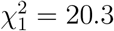, *P* < 0.001). In addition, the variance of their responses was significantly more important in open-loop condition than in closed-loop condition (Levene test on the variance: *F* = 22.6, *P* < 0.001) Box plots represent the interquartile range with the median at their center. The white dots and lines represent the means surrounded by their standard errors. Each point corresponds to the response of an individual insect exposed to an external rotation toward the right or the left. The lines connect the responses of the same individual exposed to a rotation in the same direction. *** = *P* < 0.001.

**Fig. S4:**
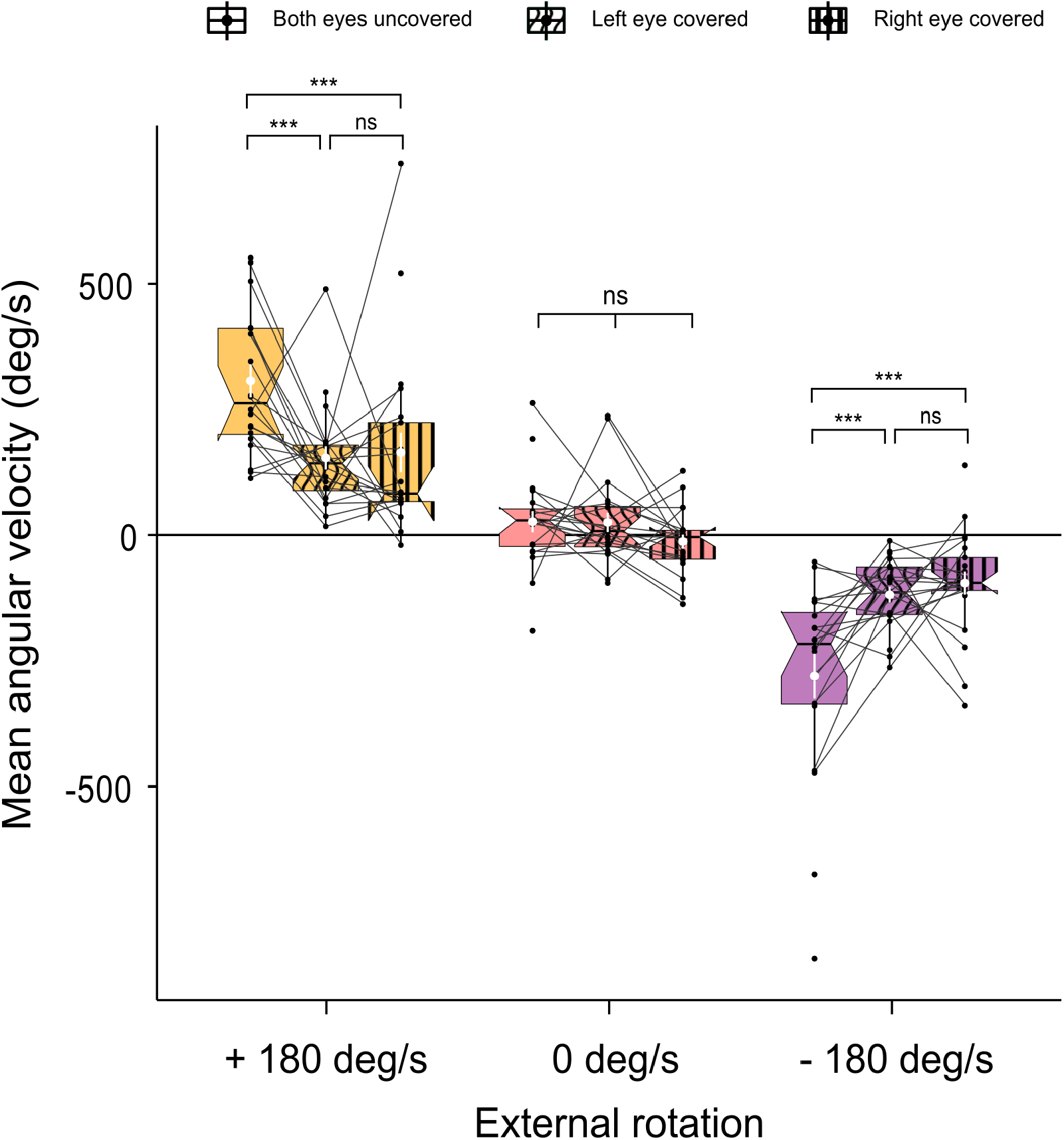
The optomotor response of monocular ants is weaker but not affected by the side of their eye. The ant’s mean angular velocity (i.e., directional bias) according to whether their right or left eye was covered with paint, and whether they were exposed or not to a constant rotating panorama toward the right (−180 deg/s) or the left (+180 deg/s). Box plots represent the interquartile range with the median at their center. The white dots and lines represent the means surrounded by their standard errors. Each point corresponds to the response of an individual ant while the lines connect the responses of the same ant across the different conditions. Data based on 22 ants tested in the VR (n=22 tested in all three conditions). We used a LMM to test for the effect of the external rotation in open-loop (factor with 3 levels: toward the right, toward the left or no external rotation), the state of their visual system (factor with 3 levels: right eye covered, left eye covered or both eyes uncovered) and the interaction between both, on the ants’ mean angular velocity. The identity of the individual was included as a random factor to account for repeated measurements. There was a significant interaction between both factors (LMM; 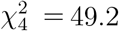 *P* < 0.001). The effect of the state of the visual system was tested for each visual condition; this effect was not significant when exposed to a fixed panorama (LMM; 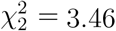, *P* = 0.177) but was significant in both optomotor conditions (rotation toward the right: LMM; 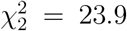, *P* < 0.001, rotation toward the left: LMM; 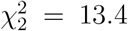, *P* = 0.001). Pairwise comparisons were done inside each optomotor group using the sequential Bonferroni correction after Holm [68]. ns = non-significant, *** = *P* < 0.001.

